# mTOR pathway gene knockout results in mTOR-dependent cellular aggregation

**DOI:** 10.1101/2025.11.02.685388

**Authors:** Kelley M. Roark, Peter B. Crino, Philip H. Iffland

**Author notes:** **Correspondence:** Philip H. Iffland II, Ph.D.

## Abstract

Malformations of cortical development (MCD) caused by variants in mTOR pathway genes (MPGs) are a leading cause of drug-resistant epilepsy. Characteristic histopathological features of MPG-associated MCD include cytomegaly and cortical dyslamination often with neurons in abnormally close apposition (aggregates). We hypothesized that cellular aggregation is an mTOR-dependent phenotype. *Tsc2, Nprl3, Stradα*, or *Kptn* were knocked out (KO) using CRISPR/Cas9 in N2a cells *in vitro*. Levels of phosphorylated ribosomal S6 protein (PS6; Ser240/244), a marker for mTOR activation, were defined via Western blotting *in vitro*. Timelapse live-cell imaging was used to observe aggregate formation, with or without mTORC1 inhibition (rapamycin). EdU-base cell proliferation assay and cell death assays were performed to determine whether aggregation was the result of changes in cell cycle or increased cell death. Liquid chromatography-mass spectrometry (LC-MS/MS) was used to define changes in the cell aggregate proteome. Human MCD brain tissue specimens were stained with PS6 to assay mTOR signaling in neuronal clusters. All knockout lines formed multi-cell aggregates compared to control lines within 24-48 hours of plating *in vitro*. Aggregation was abolished with mTOR inhibitor treatment, establishing the mTOR-dependency of aggregate formation. Aggregation was not driven by cell proliferation, apoptosis/necrosis, or the presence of extracellular DNA in culture media. LC-MS/MS analysis revealed altered expression of protein across KO lines including adhesion molecules (e.g., contactin-3), cytoskeletal proteins (e.g., stathmin-2), and protein processing/transport (e.g., Uevld). Our findings establish aberrant cellular aggregation as an mTOR-dependent phenotype across multiple MPG associated with MCD. Changes in expression of adhesion molecules may contribute to abnormal cell aggregation and cortical lamination in MCD and results in abnormal network formation that leads to seizures.

**Highlights:**

- In human MCD specimens, neurons are frequently observed in clustered groups.
- *In vitro* models of mTORopathies show mTOR-dependent changes in cellular aggregation.
- Proteomic analysis revealed changes in protein levels in adhesion molecules and other molecules relevant to cellular dynamics and protein transport.

## Introduction

Malformations of cortical development (MCD) are neurodevelopmental disorders defined by abnormal cortical cytoarchitecture and clinical association with epilepsy, intellectual disability, and autism spectrum disorder ^1,2^. Histologically, MCDs are variably defined by cortical dyslamination, cytomegalic dysmorphic neurons, heterotopic neurons in white matter, and in specific contexts, balloon cells ^3,4^. Germline or somatic loss-of-function mutations in MPGs that negatively regulate mTORC1 are increasingly recognized as key etiological drivers in MCD leading to their unification under the umbrella term “mTORopathies” ^5–7^ caused by variants in *TSC1* and *TSC2*, components of the GATOR1 complex (*NPRL2, NPRL3*, DEPDC5), the KICSTOR complex, and the pseudokinase *STRADα* ^8–10^.

While the phenotypic manifestations of MPG-associated MCD can vary by gene and mosaic variant burden, studies in human tissue consistently reveal enlarged neurons within dysplastic regions in close apposition ^11^. While mTORC1 has a known role in cell hypertrophy and altered migration, a link to neuronal aggregation has been suggested in mouse models. For example, in astrocyte-specific *Tsc2* conditional knockout mice, aberrant neuronal organization has been reported in the hippocampus with rings of neurons seen in the CA subfields ^12^. Similarly, murine *Tsc2* deletion in radial glial progenitors yields disrupted cortical cytoarchitecture and abnormal proximity between cortical neurons ^13^. Histopathological studies of cortical tubers show elevated expression of cell adhesion molecules (CAMs) like ICAM1, CD44, VCAM1, and contactin-3 in regions of neuronal clustering and balloon-cell-rich regions-pointing towards a mechanistic role of adhesion molecules in cellular aggregation ^14–16^. These converging lines of evidence suggest cellular aggregation may be a recurrent histopathological feature linked to mTOR pathway hyperactivation during brain development.

We investigated cellular aggregation as an mTORC1-dependent phenotype in the setting of MPG knockout (KO) *in vitro*. We hypothesized that MPG KO would result in mTOR-dependent aggregation and changes in adhesion molecule expression. Using CRISPR/Cas9-gene editing in N2a cells, live-cell imaging, immunochemistry, and proteomics, we identify consistent aggregation across *Nprl3, Stradα, Kptn*, and *Tsc2* knockout lines that was prevented with mTOR inhibition. Aggregation was accompanied by dysregulation of adhesion molecules, cytoskeletal components, and protein transport within proteomic datasets. These results suggest abnormal cell aggregation as a shared downstream phenotype of mTORC1 hyperactivation with potential relevance to cortical dyslamination in MPG-related MCD.

## Materials and Methods

### Assessment of neuronal aggregates in human brain tissue specimens

Human brain tissue specimens were obtained from individuals who underwent surgical resection of an epileptogenic lesion or from *post-mortem* tissue specimens. Genetic variants in brain specimens were ascertained through clinical diagnostics during routine patient care. Specimens were fixed in 4% paraformaldehyde and paraffin embedded. Using a microtome, 10 _μ_m sections were mounted on slides, dried overnight, and deparaffinized prior to staining. Brain specimens were probed for phospho-S6 (235/236) overnight at 4???????????????????C followed by incubation with biotinylated secondary antibodies (1:1000; Vector Labs) for 2 hrs. at room temperature. Cells were visualized using the VECTASTAIN ABC standard kit per the manufacturer’s protocol (Vector Labs). Slides were coverslipped with Permount (ThermoFisher) and imaged on a Leica brightfield microscope.

### Generation and validation of CRISPR/Cas9 knockout cell lines

Mouse neuro2a cells (N2aC; ATCC CCL-131) were cultured in Dulbecco’s Modified Eagle Medium (DMEM; ThermoFisher) supplemented with 10% fetal bovine serum (FBS; MilliporeSigma) under standard conditions (37 °C, 5% CO_2_). CRISPR/Cas9-mediated KO of *Tsc2, Stradα, Kptn*, or *Nprl3* was achieved using *in silico*-designed single guide RNAs (sgRNAs) cloned into the pX330-U6-Chimeric_BB-CBh-hSpCas9 vector (Addgene #42230) containing a fluorescent reporter (mCherry) using Lipofectamine LTX with Plus reagent (ThermoFisher) per the manufacturer’s protocol. sgRNAs were GCTGCAGGACAGTAGCAATT (*Nprl3*^17^), TTCTCTGGGACCACAGACGGCGG *(Tsc2*^18^), AGTCGCCATTGGAAGGCCGGAGG (*Stradα*^19^), AACAAGTCACCCCCAAAACGGGG (*Kptn)*, and GACTACCAGAGCTAACTCA (scramble control^17,18^). Forty-eight hrs. post-transfection, cells were FAC-sorted by reporter-positive expression using a BD FACS Aria II and expanded. Gene editing was verified by PCR amplification of the edited region followed by next-generation sequencing of PCR amplicons (Mass General CRISPR core). Alignment tables were generated based on WT control samples and assessed for high degrees of misalignment between experimental and control sequences.

RT-qPCR was used to define changes in mRNA expression after KO using three biological replicates for each line. Total RNA was extracted from cell pellets using the RNeasy Mini Kit (Qiagen) and RNA was converted to cDNA using a cDNA reverse transcription kit (Qiagen). Mouse GAPDH was used as an expression control (F= AGGTCGGTGTGAACGGATTTG; R= TGTAGACCATGTAGTTGAGGTCA). Using SYBR Power Green PCR master mix (ThermoFisher), a standard dilution was created using WT cDNA at the following concentrations: 1:20, 1:60, 1:180, 1:540, 1:1620. Negative controls used were DNase free water and diluted random primers. Each sample was loaded into a MicroAmp optical 384-well reaction plate at 1:20 and topped with a MicroAmp adhesive film. Plates were assayed using Viia7 qPCR machine. Analysis was performed as previously published^20^.

### Western blot assay

For protein level validation of KO lines and assessment of changes in mTOR signaling, whole-cell lysates were created using RIPA buffer containing protease and phosphatase inhibitors (Roche). Protein concentration was quantified using a NanoDrop spectrophotometer (ThermoFisher). Thirty micrograms of protein were resolved on 4-14% NuPage gels, transferred onto PVDF membranes (MilliporeSigma), blocked in Odyssey blocking solution (Li-Cor Environmental) and probed overnight at 4???????????????????C with primary antibodies against phospho-S6 (Ser240/244; Cell Signaling Technology; 1:1000), total S6 (Cell Signaling Technology; 1:1000), and β-actin (MilliporeSigma; 1:10,000) diluted in Odyssey antibody diluent (Li-Cor Environmental). To visualize primary antibody staining, Odyssey IRDye secondary antibodies were used (Li-Cor environmental; 1:1000) for 2 hrs. at room temperature. Membranes were imaged on an Odyssey CLx Scanner (Li-Cor Environmental). All Western blots were run in triplicate and quantified by densitometry in FIJI^21^ and statistically analyzed using a one-way ANOVA with Tukey’s post-hoc test.

### Imaging Cytometry

Cell size measurements were performed using the Amnis FlowSight Imaging Cytometer (Cytek Bioscience) using live cells. N2aC lines were grown under standard conditions in physiological media. Cells were trypsinized (0.05%), centrifuged at 1,500 RPM to remove physiological media resuspended in PBS containing 5% BSA, passed through a cell strainer to create a single-cell suspension, and processed on the imaging cytometer. Image segmentation and soma area quantification were performed using the accompanying software, with nuclear gating thresholds applied to exclude overlapping cells. Each condition was run in triplicate. Data were exported to Prism (Graphpad) for statistical analysis. Statistical analysis was performed using one-way ANOVA with Brown-Forsythe and Welch’s correction with a p < 0.05 deemed significant.

### Time-lapse live imaging of cellular aggregation

N2aC lines were plated in 35 mm MatTek glass-bottom dishes at a density of 2.5 × 10^5^ cells per dish and allowed to adhere for 24 hrs. Cultures were transferred to an on-stage incubator on an Olympus VivaView. Time-lapse imaging was conducted using a 10× objective over a 48-hr. period with images acquired every 15 minutes in three randomly selected regions per dish. Parallel experiments were performed in the presence of rapamycin (50 nM) added at the start of the recording. Image sequences were compiled using FIJI^21^.

### Cell proliferation assay

EdU incorporation assay was performed using the Click-iT EdU Alexa Fluor 647 Imaging Kit (ThermoFisher) according to the manufacturer’s protocol. N2aC were plated at equal density in chamber slides, incubated with 10 μM EdU for 2 hrs., and fixed in 4% paraformaldehyde. Nuclei were counterstained with DAPI, and EdU-positive cells were quantified from 10 random fields per condition using a 20× objective on a Nikon W1 spinning-disk confocal microscope. Cell counts were performed in FIJI^21^ and the percent of dividing cells was calculated as the number of EdU positive cells divided by total (DAPI positive) cells.

### Apoptosis and necrosis assays

Annexin V/Propidium iodide (PI) Apoptosis Detection Kit (Invitrogen) was used to identify apoptotic and/or necrotic cells, respectively. N2a cells were plated in 2-well chamber slides and maintained in standard growth conditions. At 48-hours post-plating, cells were stained with Annexin V–eFluor™ 488 and PI per the manufacturers protocol. Cells were fixed in 4% PFA, coverslipped with DAPI and imaged using a Leica fluorescent microscope. At least three random non-overlapping fields per well were acquired in each condition. A positive control was included by treating WT cells with etoposide (20 mM, 24 h). Images were analyzed using FIJI^21^ and cell death was quantified following Annexin V and propidium iodide (PI) staining. The number of Annexin V+/PI+ cells was calculated for each KO line.

### Assessment of cell-free DNA by qPCR

Cell-free DNA was measured in conditioned media using a β-globin–targeted quantitative PCR assay adapted from previously published protocols^20^. Briefly, DNA was isolated from 500 μL of culture media using the QIAamp DNA Mini Kit (Qiagen), and real-time PCR was performed using SYBR Green master mix (ThermoFisher) with primers specific to the mouse β-globin gene (F= CAGGCTCCTGGGCAATATGAT; R= AGCAGAAAAGGGGCTTAGTGG). Cycle threshold values from media were compared to genomic DNA extracted from pelleted cells to determine relative extracellular DNA abundance. Plates were assayed using Viia7 qPCR machine and analyzed as described above.

### Immunocytochemistry

Cells were plated in 2 well chamber slides and fixed at 48 hrs. post-plating using 4% paraformaldehyde (PFA) for 15 minutes at room temperature. Following fixation, cells were permeabilized in 0.1% triton X-100 in PBS for 10 minutes and then blocked in 5% normal goat serum (NGS; Vector Labs) in PBS for 1 hour. To visualize cytoskeletal architecture, cells were incubated with primary anti-F-actin antibody (1:500 in blocking buffer; Abcam) overnight at 4°C, followed by incubation with goat anti-rabbit Alex Fluor 568-conjugated secondary antibody (1:1000; ThermoFisher) for 1 hour at room temperature in the dark.

Nuclei were counterstained with DAPI (1ug/mL; ThermoFisher) and slides were mounted using Vectashield Antifade Mountant (Vector Laboratories). Images were acquired using a W1 spinning-disk confocal microscope. Aggregate number, area, and nuclei per aggregate were quantified using FIJI^21^. Aggregates were defined as dense and discrete areas of cells within a field. All experiments were performed in triplicate. Data were statistically analyzed by one-way ANOVA, with p < 0.05 considered significant.

### Mass spectrometry and proteomic analysis

To identify protein-level changes associated with cellular aggregation in MPG models, we performed quantitative proteomic profiling of the detergent-insoluble (membrane-bound/cytoskeletal-enriched) fraction from N2aC lines. Proteins were extracted using cloud point separation in 2% Triton-X114. Three biological replicates were submitted for each of the following conditions: scramble control, *Kptn* KO, *Nprl3* KO, *Stradα* KO, and *Tsc2* KO. Sample processing, peptide digestion, and liquid chromatography tandem mass spectrometry (LC-MS/MS) were performed by the University of Maryland School of Medicine Protein Analysis Lab using a ThermoFisher LTQ ion trap mass spectrometer. Data were analyzed in LFQ-Analyst (Monash University). Changes in proteins were considered statistically significant if the p < 0.001 and had a two-fold or greater change in expression.

### Study Approval

This study was approved by the University of Maryland School of Medicine Institutional Review Board and Institutional Biosafety Committee.

## Results

### Cellular Aggregation is a shared histopathological feature in human MCD specimens

Histological analysis of resected human cortical tissue from individuals with pathogenic *NPRL3* (FCD II), *TSC*, and *STRADα* variants revealed clusters of dysmorphic, cytomegalic, and PS6 positive neurons in aggregates or clusters (no brain tissue specimens from individuals with *KPTN* variants exist to assay). These structures were morphologically distinct from the radial dyslamination and balloon cells characteristic of tuberous sclerosis complex and FCD2. Indeed, each MCD exhibited disorganized aggregates of large neurons in close physical apposition with reduced intercellular spacing. These features were absent from control cortex and suggest that cellular aggregation may represent a distinct, underrecognized, histopathological hallmark of mTOR pathway gene loss-associated MCD. **(Figure 1)**.

**Figure 1.**
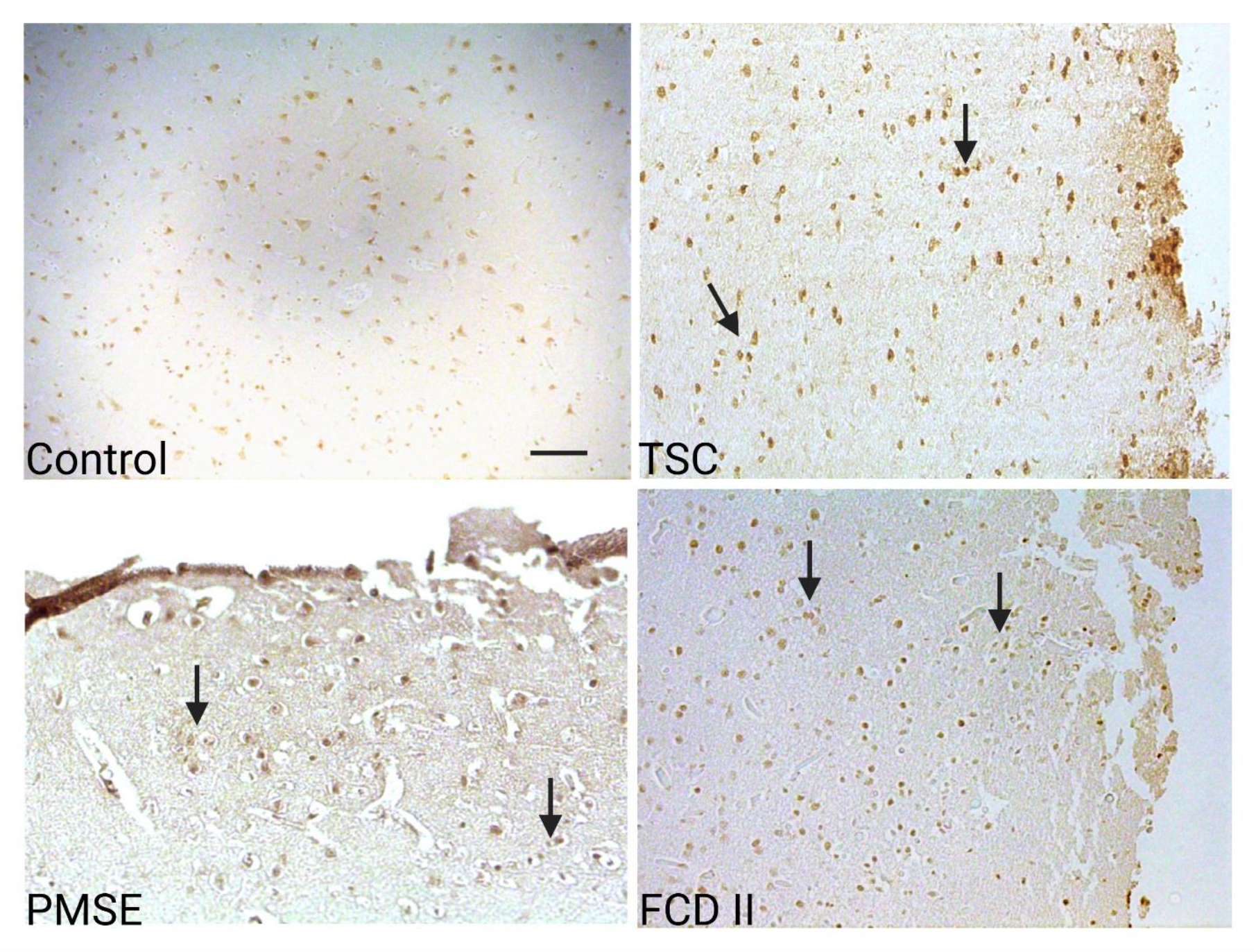
Histopathological evidence of cellular aggregation in MPG variant-associated MCD. Representative PS6-stained, or NeuN for control, cortical sections from individuals with TSC, PMSE (*STRADα)*, or FCD II (*NPRL3)*, showing clustered cytomegalic neurons with reduced intercellular spacing (*arrowheads*). Scale bars = 100 μm. TSC = tuberous sclerosis complex, PMSE = polyhydramnios megalencephaly and symptomatic epilepsy, FCD II = focal cortical dysplasia type II.

### CRISPR/Cas9-mediated KO of MPGs induces mTORC1 hyperactivation and cell hypertrophy

To model MPG knockouts, we used CRISPR/Cas9 to generate *Nprl3, Stradα, Kptn*, and *Tsc2* knockout lines in N2aC. The *Nprl3, Tsc2*, and *Stradα* KO cell lines have all been previously validated^17–19^. Cas9 editing of regions of interest was confirmed by next-generation sequencing assaying for mismatched DNA pairs within the edited region *(Data not shown or published elsewhere)*. Western blot analysis (n=3 per group) confirmed statistically significant hyperphosphorylation of ribosomal protein S6 (PS6; Ser240/244), indicating mTOR pathway hyperactivation in each KO line relative to controls. Cell size, quantified using automated imaging cytometry measuring cell area, was significantly increased in all KO lines relative to WT and scramble controls and was prevented with rapamycin treatment **(Figure 2)**. Statistical analysis was performed using one-way ANOVA with Welch’s correction (W (11, 12762) = 417.5, p < 0.0001; total n= 50,753 cells). Multiple comparisons were conducted using Welch’s t-test with multiplicity-adjusted p-values. WT and Scramble controls showed no significant change in soma size with rapamycin as we have observed previously^22^.

**Figure 2.**
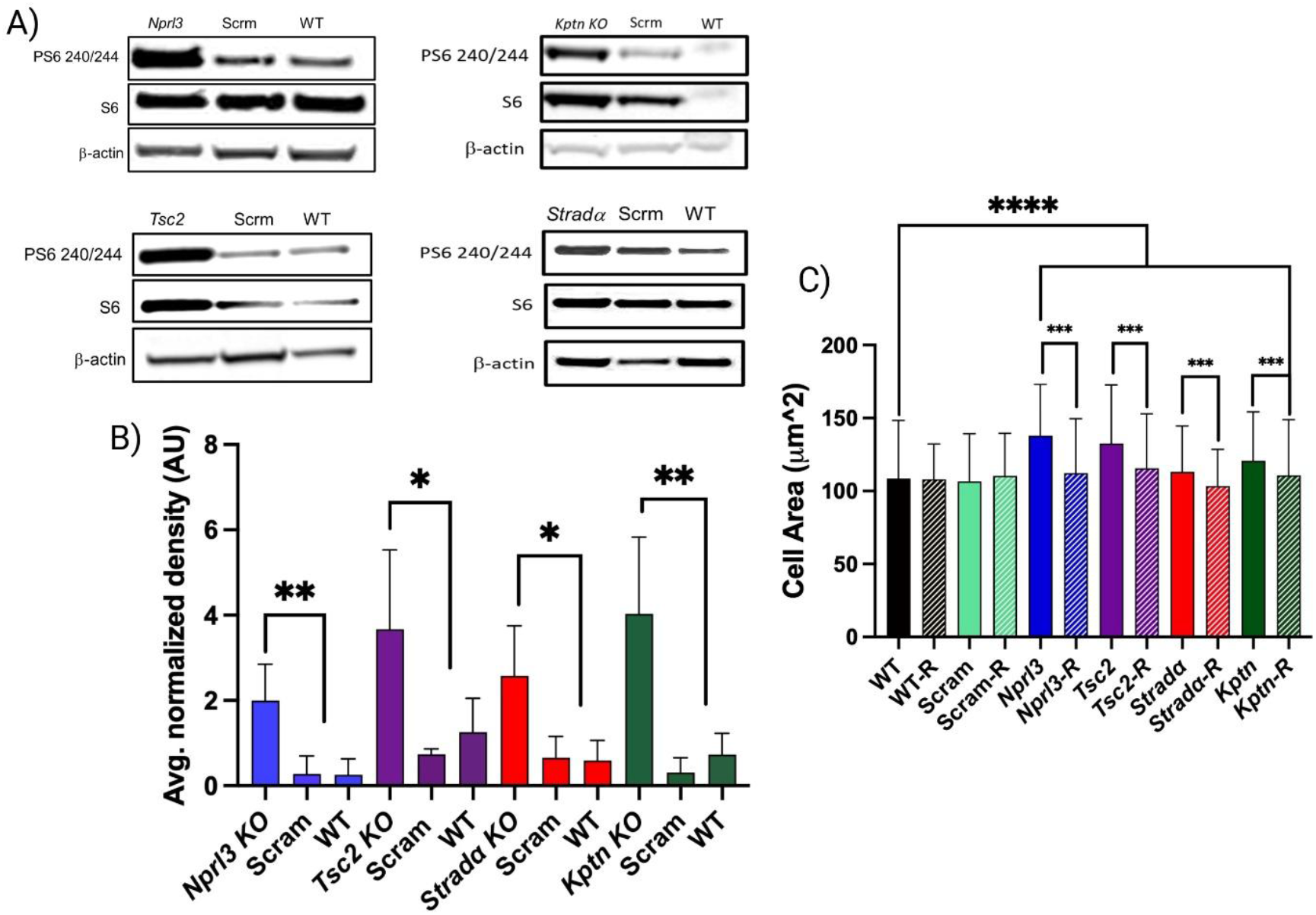
CRISPR/Ca9 knockout of MPGs induces mTORC1 hyperactivation and cell hypertrophy. **A)** Representative western blots showing PS6 (ser240/244) and total S6 levels in wildtype, scramble control, and knockout N2a lines (*Nprl3, Stradα, Kptn*, and *Tsc2*). All KO lines exhibit elevated PS6 signal relative to controls. β-actin was used as a loading control. **(B)** Quantification of PS6 band intensity normalized to β-actin (n=3). Data are shown as mean ± SD from three biological replicates. **(C)** Imaging cytometry analysis of cell soma diameter from _≥_5,000 cells per group reveals significant cellular hypertrophy in all KO lines compared to controls that was prevented with rapamycin (50 nM; 48 hrs). **** = p<0.0001, *** = p<0.001, **= p<0.01, * = p< 0.05. WT = wildtype, scram = scramble control, R = rapamycin, T= torin1.

### MPG knockout leads to mTORC1-dependent cellular aggregation and is not caused by increased proliferation or cell death

In untreated MPG KO (*Nprl3, Strada, Kptn, and Tsc2)* samples, aggregates were significantly more numerous, had greater numbers of cells, and had a larger area compared to both WT and scramble control **(Figure 3)**. Aggregate formation, as measured by aggregate number, nuclei count, and area was prevented with rapamycin or torin1 treatment. Live-cell timelapse imaging revealed that all four KO lines formed large, motile aggregates within 48 hrs. of plating. These structures grew over time, incorporating surrounding cells, and migrated collectively as dense units (**Supplemental video 1)**. No aggregation was observed in wildtype or scramble control lines. Treatment with rapamycin or torin1 prevented aggregate formation, confirming mTORC1-dependency of these changes.

**Figure 3.**
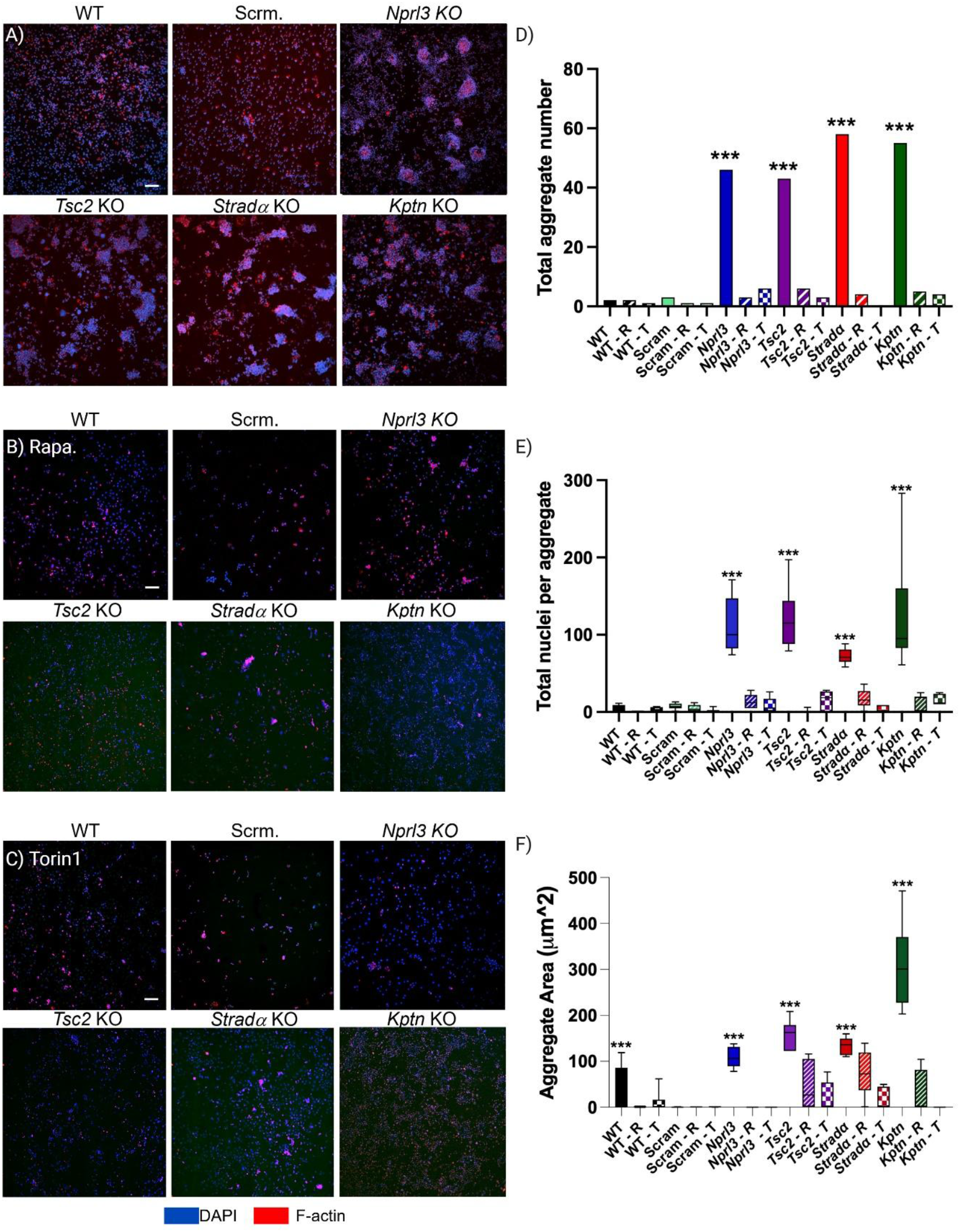
Knockout of MPGs drive mTORC1-dependent aggregation *in vitro*. **(A-C)** Representative images from of N2a cells allowed to aggregate for 48 hrs. **(A)**. In **B** and **C**, representative images of cells allowed to aggregate for 48 hrs. and incubated with rapamycin (50 nM) or torin1 (50 nM), respectively (n=3 replicates per cell line). In physiological media, large aggregates **(A)** form in each KO line compared to control as measured by total aggregate number, nuclei per aggregate, and aggregate volume **(D-F)**. mTOR inhibitor treatment resulted in statistically significant decreases in aggregate number, nuclei per aggregate, and aggregate volume across all KO lines **(B**,**C, D-F)**. Scale bar = 200 _μ_m *** = p<0.001, **= p<0.01, * = p < 0.05. WT = wildtype, scram = scramble control, R or rapa = rapamycin, T= torin1. F-actin = filamentous actin.

To determine whether cell aggregates formed as a consequence of mTOR-induced cell proliferation, an Edu-based cell proliferation assay was performed **(Figure 4)**. Cell division was quantified by determining the percentage of dividing cells (Edu positive). Across experimental groups, no changes in cell division were observed (n= approx. 300 cells per group). Apoptosis and necrosis were assayed using annexin V and propidium iodide, respectively **(Figure 4)**. No increases in either apoptosis or necrosis were observed in KO cell lines relative to an etoposide treated positive control (n= approx. 300 cells per group). In addition, cell death can trigger the release of DNA into cell culture media which can increase cell-to-cell adherence. The presence of cell-free DNA in cell culture media was assayed using qPCR probing for mouse _β_-globin and compared to intracellular _β_-globin levels using WT genomic DNA **(Figure 4)**. No statistically significant increase in cell-free DNA was observed across experimental KO cells (p > 0.05, n= approx. n=3 replicates per group).

**Figure 4.**
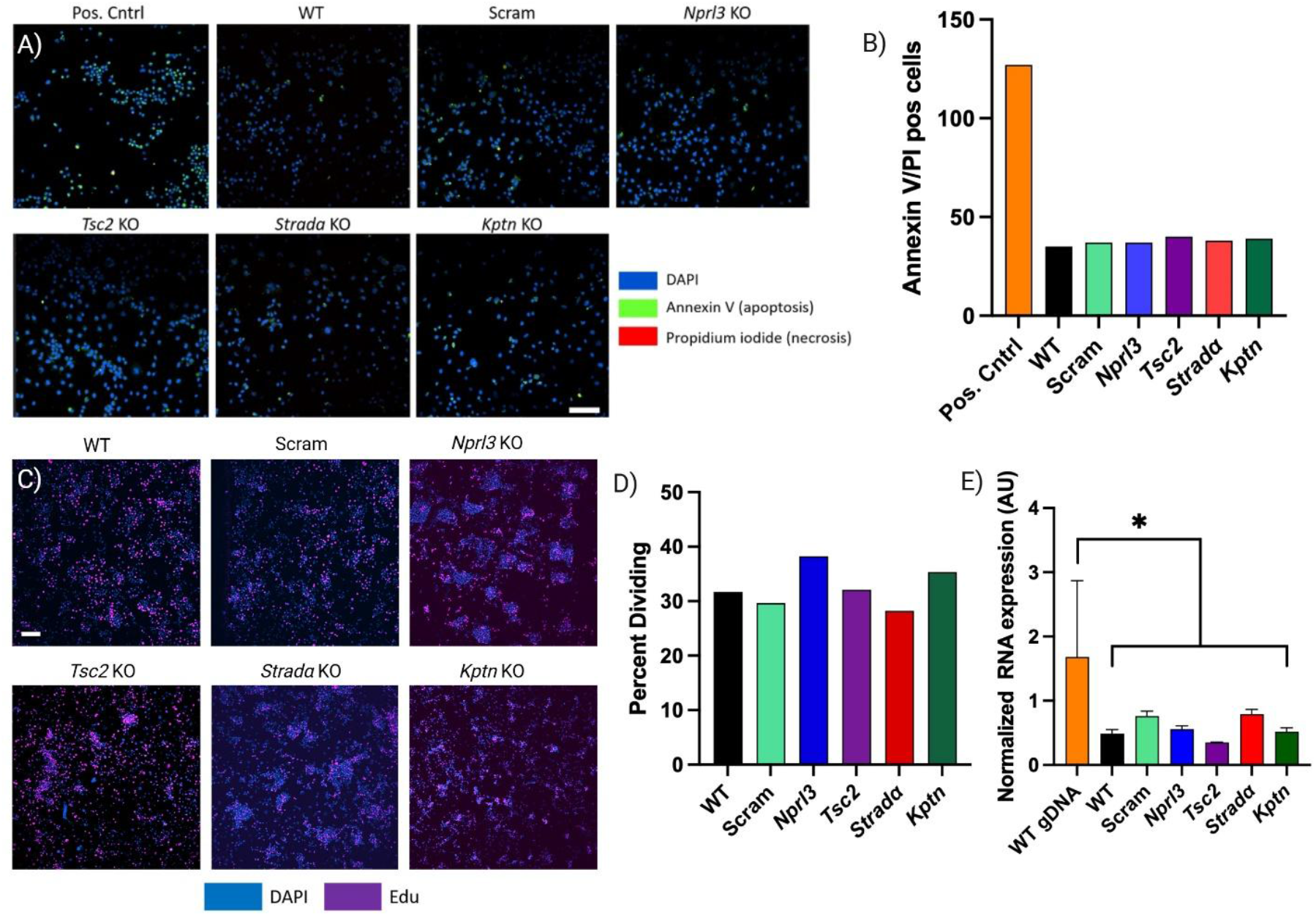
Cell aggregation is not caused by increased proliferation or cell death. **(A)** Representative images of WT, scramble, *Nprl3, Stradα, Kptn*, and *Tsc2* KO N2a cell lines subjected to Annexin V and PI staining. A positive control, WT cells incubated with etoposide, is also shown. No increase in cell death was observed relative to the positive control or WT and scramble cells **(B)**. Cell death was also assayed by measuring cell-free DNA (mouse _β_-globin) in culture media relative to genomic DNA. No statistically significant changes in levels of cell-free DNA were observed relative to WT or scramble control, but genomic _β_-globin was expectedly high **(E)**. An edu-based cell proliferation assay (**C)** revealed no changes in proliferation **(D)** across all cell lines. Scale bars: 100 μm (A), 50 μm (D). * = p < 0.05.

### Proteomic profiling reveals broad protein dysregulation in MPG KO cells

To identify the potential molecular drivers of the aggregation phenotype, we performed LC/MS proteomic profiling on detergent enriched membrane fractions from each KO line and scramble control. Quality control data for this experiment can be found in **Supplemental Figure**

**1**. Differentially expressed proteins were considered statistically significant at a p< 0.001 and a log fold change of _≥_ ±2. There were several common differentially expressed proteins across KO lines including SLC25A24 which was down-regulated in all samples. Additionally, Capn2 and Scaf8 were down-regulated and Stmn2 and Scag2 were up-regulated in *Stradα* and *Kptn* **(Figure 5**). In comparing N*prl3* and *Tsc2*, Pcsk1n was down-regulated in both, and Lsm1 and Uevld were upregulated, along with changes in other proteins **(Supplemental data 2)**. Interestingly, there were more differentially expressed proteins when comparing *Nprl3* vs. *Tsc2* than in comparing *Strada* vs. *Kptn* - where only 40 proteins were differentially expressed **(Figure 5)**.

**Figure 5:**
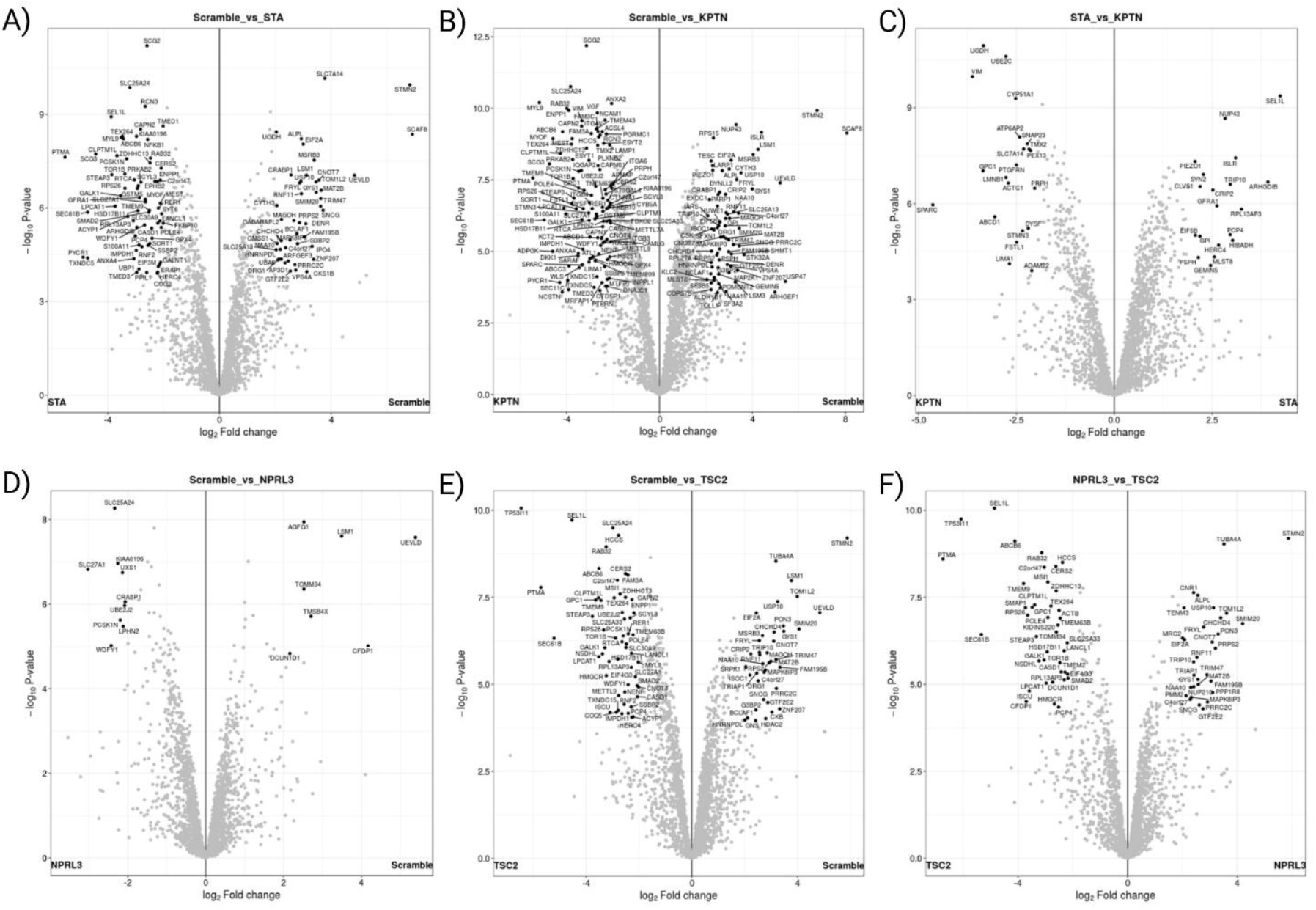
Proteomic analysis reveals broad changes in protein expression in MGP KO lines. Volcano plots showing differentially expressed proteins comparted to control are shown in **A**,**B**,**D**, and **E**. In **C** and **F**, comparison between *Strada* and *Kptn* and *Nprl3* and *Tsc2* are shown, respectively. All l*abeled* and *bolded* proteins are statistically significant at p< 0.001 and a log fold change _≥_ ± 2. SLC25A24 was downregulated in all KO lines relative to control. Capn2, Stmn2, Scaf8, and Scg2 were altered in both *Kptn* and *Stradα* KO lines **(A**,**B)**. Pcsk1n, Lsm1, and Uevld were dysregulated in both *Nprl3* and *Tsc2* KO lines **(D**,**E)**. Comparison between both *Kptn* and *Stradα* KO lines revealed relatively few differentially expressed proteins **(C)** but a greater number of proteins were expressed differently between *Nprl3* and *Tsc2* KO lines **(F)**. STA = Strad_α·_

To assay changes in cellular processes more closely, KEGG (Kyoto Encyclopedia of Genes and Genomes) revealed differential expression of multiple pathways/processes including several associated with cell adhesion. KEGG enrichments analysis showed overrepresentation of pathways related to cell-to-cell junctions, protein processing in the endoplasmic reticulum, and cytoskeletal function **(Supplemental data 1)**. Notably, in support of prior human studies, *contactin-3 (CNTN3;* **Supplemental Figure 2)**, was significantly down-regulated in *Tsc2* KO cells (but not other KO lines) and several cadherins (CDH13, CDH11), and integrins (ITGB1) were also reduced across KO lines **(Supplemental data 2)**. These findings implicate CAM, cytoskeletal, and protein processing dysregulation as a shared downstream consequence of mTORC1 hyperactivation and a potential mechanism for cellular aggregation.

## Discussion

We demonstrate that knockout of MPG (e.g., *Nprl3, Kptn, Stradα*, and *Tsc2*) induces robust, mTORC1-dependent aggregation of cells *in vitro*. The aggregation phenotype is absent in wildtype or scramble controls, independent of changes in proliferation or cell death, and is suppressed by pharmacologic inhibition of mTORC1 signaling. Aberrant cellular aggregation caused by MPG KO *in vitro* may represent a distinct form of cortical disorganization with potential relevance to MCD and epilepsy. Together, these data support our hypothesis that mTOR hyperactivation caused by MPG KO leads to abnormal cell aggregation which result from altered expression of CAMs.

Clusters of abnormally spaced neurons as well as densely packed, cytomegalic neurons are a histopathological feature of focal cortical dysplasia type 2, tuberous sclerosis complex, and hemimegalencephaly^11^. We submit that cellular aggregation *in vitro* reinforces the relevance of this phenotype and raises the possibility that pathological aggregation may contribute to the disrupted architecture seen in cortical malformations ^17^. Our findings suggest that altered cell-to-cell adhesion may represent an additional axis of developmental disorganization.

Mechanistically, our data point toward dysregulation of cell adhesion molecules (CAMs) as a plausible driver of aggregation. Proteomic profiling revealed consistent changes in multiple CAM families—including contactins, cadherins, and integrins—across knockout lines. Notably, Contactin-3 (CNTN3), a synaptic adhesion molecule implicated in neuronal spacing and arborization, was significantly downregulated in *Tsc2* KO cells and is supported by recent transcriptomic evidence showing selective downregulation of CNTN3 in cortical tubers from TSC patients. Our findings implicate CNTN3 as a candidate adhesion-related vulnerability in mTOR pathway associated MCD.

Although further validation is required, these findings support a model in which mTORC1 hyperactivation disrupts adhesive equilibrium, allowing for inappropriate cell–cell cohesion during early neuronal development. Indeed, mTORC1 activity directly modulates adhesion by controlling integrin trafficking, cytoskeletal remodeling, and translation of adhesion-associated transcripts via S6K and 4E-BP1 signaling ^23,24^. Thus, mTORC1 hyperactivation may induce a cellular state in which increased adhesiveness drives clump formation and impedes normal spacing between neurons – a possibility supported by histological findings in TSC, where CAMs such as CD44 and VCAM1 are enriched in regions of balloon cell clustering ^14^.

By highlighting a shared cell-intrinsic, adhesion-related vulnerability among mTORopathies driven by mTORC1 hyperactivation, this work expands current models of cortical disorganization and introduces a novel axis of pathology with potential relevance to epileptogenesis. Future studies aimed at defining the structural, functional, and developmental consequences of this aggregation phenotype will be critical for understanding how altered cell-to-cell interactions contribute to the pathogenesis of malformations of cortical development.

We acknowledge several limitations to our study. First, immortalized N2a neuroblastoma cells, though advantageous for controlled genetic manipulation and live-cell imaging, provide an incomplete representation of developing cortical neurons within the *in vivo* neuroepithelium. Future work employing primary neuronal cultures, brain organoids, or *in vivo* mosaic models will be needed to determine whether similar mTOR-dependent aggregation occurs within cortical laminae. Second, the mechanistic connection between adhesion molecule dysregulation and aggregate formation also remains correlative. Although the data implicate cadherins, integrins, and contactins as candidate mediators, direct functional studies will be essential to establish causality and determine whether specific adhesion pathways are sufficient or necessary for the aggregation phenotype.

Finally, while the presence of PS6-positive clusters in human MCD tissue supports translational relevance, interpretation is limited by sample heterogeneity and small cohort size. The specimens varied in genotype, cortical region, and developmental stage, and the lack of - tissue from KPTN-related disorder restricts cross-species phenotypic comparison for this genotype. Broader histopathological and molecular analyses across genotypically defined cohorts will be critical to clarify whether neuronal clustering represents a common or gene-specific feature within the mTORopathy spectrum.

## Acknowledgements

The authors would like to thank Mr. Brian Hampton for proteomic services. All figures created in https://BioRender.com

## Author contributions

KMR: formal analysis, investigation, methodology, validation, visualization, writing-original draft, writing-review and editing. PBC: conceptualization, funding acquisition, resources, supervision, writing-review and editing. PHI: conceptualization, data curation, formal analysis, funding acquisition, investigation, methodology, project administration, resources, supervision, writing-original draft, writing-review and editing.

## Figures and figure legends

**Supplemental Figure 1:**
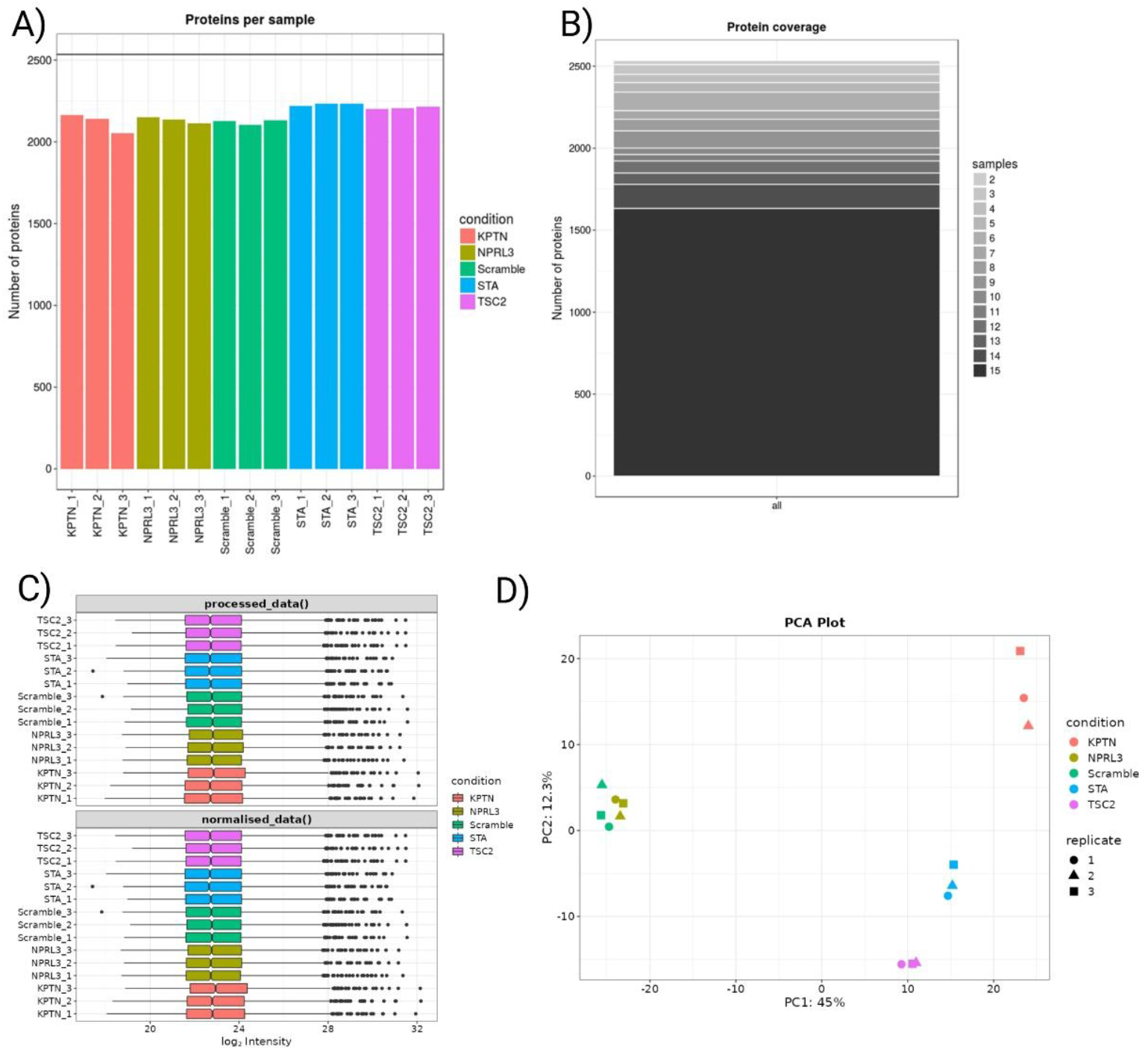
Proteomics quality control data. An approximately equal number of proteins were samples across each replicated in each KO cell line **(A)**. Additionally, a high number of possible proteins were covered within the data set **(B)** and were normalized across KO lines and replicates **(C)**. Lastly, principal components analysis revealed that all KO lines were different from control with *Nprl3* KO being the least different and *KPTN* being the most different. Interestingly, *Tsc2* KO was more similar to *Stradα* KO than *Kptn* KO.

**Supplemental Figure 2:**
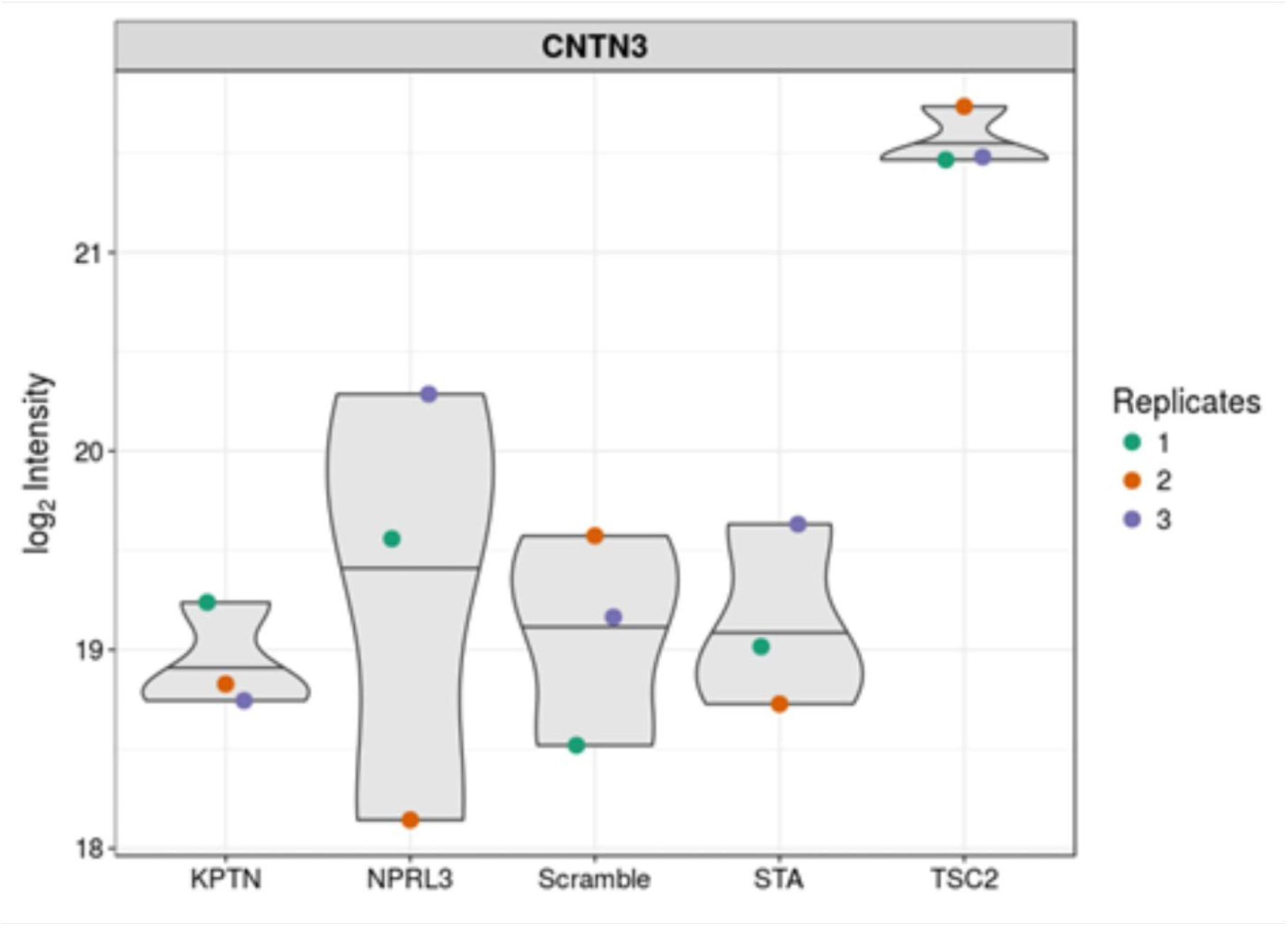
Contacting-3 levels are decreased only in *Tsc2* KO cells. Contactin-3 was previously shown to be down-regulated in humans TSC brain specimens^16^. Our proteomics data also revealed a statistically significant (adj p = 0.00108) decrease. Contactin-3 was specifically down-regulated compared to scramble control and all other MGP KO lines. Note, in this graph, higher levels on the graph indicated a decrease.

**Supplemental video: Timelapse videos of cellular aggregation across MPG KO lines with or without mTOR inhibitor treatment**. A 45 second video showing timelapse imaging of cellular aggregation in real-time. Images were acquired every 15 minutes for 48 hrs. In the first 15 seconds, baseline videos are shown of all KO and control lines. Large aggregates were observed in all KO lines. In the second and third 15 second segments, videos show cells incubated with rapamycin (50 nM, 48 hrs.) or torin1 (50 nM; 48 hrs), respectively. No aggregates, or only a few small static aggregates, were observed in mTOR inhibitor treated cell lines.

**Supplemental data 1: KEGG enrichment analysis data table**. Complete data set of altered pathways found within all cell lines and in comparison to all other samples.

**Supplemental data 2: Proteomics data files:** proteomics data can be access by uploading the files to https://analyst-suite.monash-proteomics.cloud.edu.au/apps/lfq-analyst//

## References

1 Aronica E, Becker AJ, Spreafico R. Malformations of Cortical Development. Brain Pathology 2012;22:380–401. 10.1111/j.1750-3639.2012.00581.x.

2 Mühlebner A, Bongaarts A, Sarnat HB, Scholl T, Aronica E. New insights into a spectrum of developmental malformations related to mTOR dysregulations: challenges and perspectives. Journal of Anatomy 2019;235:521–42. 10.1111/joa.12956.

3 Blumcke I, Budday S, Poduri A, Lal D, Kobow K, Baulac S. Neocortical development and epilepsy: insights from focal cortical dysplasia and brain tumours. The Lancet Neurology 2021;20:943–55. 10.1016/S1474-4422(21)00265-9.

4 Crino PB. mTOR Signaling in Epilepsy: Insights from Malformations of Cortical Development. Cold Spring Harb Perspect Med 2015;5:a022442. 10.1101/cshperspect.a022442.

5 Moller RS, Weckhuysen S, Chipaux M, Marsan E, Taly V, Bebin EM, et al. Germline and somatic mutations in the MTOR gene in focal cortical dysplasia and epilepsy. Neurol Genet 2016;2:e118. 10.1212/NXG.0000000000000118.

6 Moloney PB, Cavalleri GL, Delanty N. Epilepsy in the mTORopathies: opportunities for precision medicine. Brain Communications 2021;3:fcab222. 10.1093/braincomms/fcab222.

7 Nguyen LH, Bordey A. Convergent and Divergent Mechanisms of Epileptogenesis in mTORopathies. Front Neuroanat 2021;15:. 10.3389/fnana.2021.664695.

8 Baple EL, Maroofian R, Chioza BA, Izadi M, Cross HE, Al-Turki S, et al. Mutations in KPTN Cause Macrocephaly, Neurodevelopmental Delay, and Seizures. Am J Hum Genet 2014;94:87–94. 10.1016/j.ajhg.2013.10.001.

9 Feliciano DM, Bordey A. TSC-mTORC1 Pathway in Postnatal V-SVZ Neurodevelopment. Biomolecules 2025;15:573. 10.3390/biom15040573.

10 Ricos MG, Hodgson BL, Pippucci T, Saidin A, Ong YS, Heron SE, et al. Mutations in the mammalian target of rapamycin pathway regulators NPRL2 and NPRL3 cause focal epilepsy. Annals of Neurology 2016;79:120–31. 10.1002/ana.24547.

11 Iffland PH, Crino PB. Focal Cortical Dysplasia: Gene Mutations, Cell Signaling, and Therapeutic Implications. Annu Rev Pathol 2017;12:547–71. 10.1146/annurev-pathol-052016-100138.

12 Uhlmann EJ, Wong M, Baldwin RL, Bajenaru ML, Onda H, Kwiatkowski DJ, et al. Astrocyte-specific TSC1 conditional knockout mice exhibit abnormal neuronal organization and seizures. Ann Neurol 2002;52:285–96. 10.1002/ana.10283.

13 Way SW, McKenna J, Mietzsch U, Reith RM, Wu HC, Gambello MJ. Loss of Tsc2 in radial glia models the brain pathology of tuberous sclerosis complex in the mouse. Hum Mol Genet 2009;18:1252–65. 10.1093/hmg/ddp025.

14 Arai Y, Takashima S, Becker LE. CD44 Expression in Tuberous Sclerosis. Pathobiology 2000;68:87–92. 10.1159/000028118.

15 Boer K, Crino PB, Gorter JA, Nellist M, Jansen FE, Spliet WG, et al. Gene expression analysis of tuberous sclerosis complex cortical tubers reveals increased expression of adhesion and inflammatory factors. Brain Pathol 2010;20:704–19. 10.1111/j.1750-3639.2009.00341.x.

16 Korotkov A, Luinenburg MJ, Romagnolo A, Zimmer TS, van Scheppingen J, Bongaarts A, et al. Down-regulation of the brain-specific cell-adhesion molecule contactin-3 in tuberous sclerosis complex during the early postnatal period. J Neurodev Disord 2022;14:8. 10.1186/s11689-022-09416-2.

17 Iffland PH II, Everett ME, Cobb-Pitstick KM, Bowser LE, Barnes AE, Babus JK, et al. NPRL3 loss alters neuronal morphology, mTOR localization, cortical lamination and seizure threshold. Brain 2022:awac044. 10.1093/brain/awac044.

18 Iffland PH, Barnes AE, Baybis M, Crino PB. Dynamic analysis of 4E-BP1 phosphorylation in neurons with Tsc2 or Depdc5 knockout. Exp Neurol 2020;334:113432. 10.1016/j.expneurol.2020.113432.

19 Dang LT, Glanowska KM, Iffland Ii PH, Barnes AE, Baybis M, Liu Y, et al. Multimodal Analysis of STRADA Function in Brain Development. Front Cell Neurosci 2020;14:122. 10.3389/fncel.2020.00122.

20 Nugent BM, Wright CL, Shetty AC, Hodes GE, Lenz KM, Mahurkar A, et al. Brain feminization requires active repression of masculinization via DNA methylation. Nat Neurosci 2015;18:690–7. 10.1038/nn.3988.

21 Schneider CA, Rasband WS, Eliceiri KW. NIH Image to ImageJ: 25 years of image analysis. Nat Methods 2012;9:671–5.

22 Iffland PH, Baybis M, Barnes AE, Leventer RJ, Lockhart PJ, Crino PB. DEPDC5 and NPRL3 modulate cell size, filopodial outgrowth, and localization of mTOR in neural progenitor cells and neurons. Neurobiol Dis 2018;114:184–93. 10.1016/j.nbd.2018.02.013.

23 Chen L, Xu B, Liu L, Liu C, Luo Y, Chen X, et al. Both mTORC1 and mTORC2 are involved in the regulation of cell adhesion. Oncotarget 2015;6:7136–50. 10.18632/oncotarget.3044.

24 Gulhati P, Bowen KA, Liu J, Stevens PD, Rychahou PG, Chen M, et al. mTORC1 and mTORC2 Regulate EMT, Motility, and Metastasis of Colorectal Cancer via RhoA and Rac1 Signaling Pathways. Cancer Research 2011;71:3246–56. 10.1158/0008-5472.CAN-10-4058.

